# Serum-free culture system for spontaneous human mesenchymal stem cell spheroids formation

**DOI:** 10.1101/666313

**Authors:** Guoyi Dong, Shengpeng Wang, Yuping Ge, Qiuting Deng, Qi Cao, Quanlei Wang, Zhouchun Shang, Wenjie OuYang, Jing Li, Chao Liu, Jie Tang, Weihua Zhao, Ying Gu

## Abstract

Human mesenchymal stem cells (hMSCs) are widely used in clinical research because of their multipotential, immunomodulatory, and reparative properties. Previous studies determined that hMSC spheroids from three-dimensional (3D) culture possess higher therapeutic efficacy than conventional hMSCs from monolayer (2D) culture. To date, various 3D culture methods have been developed to form hMSC spheroids, but most of them used culture medium containing fetal bovine serum (FBS), which is not suitable for further clinical use. Here, we demonstrate that dissociated single MSCs seeded in induced pluripotent stems medium (MiPS), adhere loosely to the dish and spontaneously migrate to form spheroids during day 3 to day 6. Through component deletion screening and complementation experiments, the knockout serum replacement (KSR) was identified as necessary and sufficient for hMSC spheroid formation. Transcriptome analysis showed that the overall expression profiles were highly similar between 2D culture with FBS and KSR derived spheroids. Interestingly, genes related to inflammatory response, immune response, and angiogenesis were up-regulated in spheroids at day 6, and qPCR results further validated the increased expression level of related genes, including *STC1, CCL7, HGF, IL24*, and *TGFB3*. When spheroids were re-plated in normal FBS medium, cells formed a typical spindle-shaped morphology, and FACS results showed that the recovered cells retained MSC-specific surface markers, such as CD73, CD90, and CD105. In summary, we developed a practical and convenient method to generate hMSC spheroids for clinical research and therapy.

## Introduction

Human mesenchymal stem cells (hMSCs) possess self-renewal and multilineage differentiation potential [1, 2] and are extensively used in clinical studies. hMSCs can be derived from a wide range of tissues, such as bone marrow, adipose tissue, umbilical cord, placenta and dental pulp [3], and can be cultured *in vitro* for several generations. However, clinical data has shown a low survival rate for hMSCs from monolayer two-dimensional (2D) culture when implanted *in vivo*, and optimized approaches for hMSCs production are required for clinical application. Recent studies showed that aggregating hMSCs into 3D spheroids increased cell survival [4], stemness [5, 6], anti-inflammatory [7] and proangiogenic [8-10] properties of hMSCs. These data imply that 3D spheroids can be an alternative source for hMSCs in clinical applications.

A variety of *in vitro* 3D spheroid culture approaches have been developed [11-13], including hanging drop [7, 14-16], pre-coating of low-adhesive substrates [17], membrane-based aggregation [5, 18, 19] and forced aggregation [20]. However, most of these methods use conditioned medium containing fetal bovine serum (FBS), which contains undefined components and is not recommended for clinical applications [21, 22]. So far, several studies using a serum-free medium have successfully generated characterized hMSC spheroids. The Yloslato group utilized various serum-free and chemically defined xeno-free media, including MSCGM, MesenCult XF, and StemPro XF to generate hMSC spheroids in hanging drops, and found that compact spheroids formed when human serum albumin (HSA) was added into MesenCult XF and StemPro XF medium. Furthermore, they demonstrated that these hMSC spheroids were activated to express higher levels of therapeutic genes, such as *TSG6, IL1A, IL1B*, and *STC1*, in StemPro XF medium supplemented with HSA [23]. Meanwhile, Zimmermann et al. utilized the forced aggregation method to form hMSC spheroids in agarose microwells and then cultured them in MesenCult XF medium, and they found that hMSC spheroids only grow in size, but did not increase the production of immunomodulatory paracrine factor PGE2 and IL-6 and IDO [20]. These studies suggested that hMSC spheroids derived from serum-free medium culture maintained, if not enhanced, the hMSCs properties; however, the complicated procedures (hanging drops or gel coating) or instrument (agarose microwells) requirements seriously hinder their potential large-scale applications.

To solve these problems, we developed a novel method to generate hMSC spheroids spontaneously in serum-free condition medium containing knockout serum replacement (KSR). RNA-seq and qPCR results showed that MSC spheroids showed up-regulated expression of therapeutic factors, including inflammatory response, immune response, and angiogenesis genes, and spheroid cells retained MSC-specific immunophenotypic markers after re-plating in FBS culture medium. Overall, our approach provides a convenient and cost-effective method to generate hMSC spheroids with therapetical potentials.

## Methods and materials

### Cell culture

hMSCs were isolated from umbilical cord tissue and cultured in L-DMEM (Gibco, 11885-084) medium containing 15% FBS (Hyclone, sh30084.03) at 37°C in a humidified atmosphere of 5% CO2, and meium were changed every three days. Cells were dissociated with 0.05% Trypsin-EDTA (Invitrogen, 25300062) and passaged at 1:3 ratio when reaching about 80% confluence. Cells from passage 3 to passage 8 were used in this study.

### Formation of spheroids and spheroid recovered monolayer MSCs

Cells at passage 3, 5 and 8 were dissociated into single cells with 0.05% Trypsin-EDTA (Invitrogen, 25300062) and seeded at the indicated concentration (1×10^4^, 2.5×10^4^, 5×10^4^, 1×10^5^ per cm^2^) in human induced pluripotent stem cells medium (refer to as MiPS), which is DMEM/F12 (Gibco, 11320-033) containing 20%KSR (Gibco, 10828-028), 2uM L-Glutamine (Sigma G8540), 0.1uM NEAA (Gibco, 11140-050), 0.1uM 2-Mercaptoethanol (Gibco, 21985-023) and 10ng/ml human bFGF (Invitrogen, PHG0021) or in L-DMEM (Gibico, 11885-084) containing 20% KSR (refer to as L-KSR). Other mediums containing 20% KSR used in this study include H-DMEM (LIFE TECHNOLOGIES, 11965-092), DMEM/F12 (LIFE TECHNOLOGIES, 10565-018), MEM (GIBCO, 11095080), RPMI1640 (GIBCO, C22400500BT). Cells were incubated at 37°C in a humidified atmosphere of 5% CO2. To recover monolayer hMSCs from spheroids, spheroids at day 6 in MiPS or L-KSR were collected and washed with PBS, then cultured in L-FBS.

### Image and Video record

Cells or spheroids were photographed with a microscope (Zeiss LSM 510) at day 1, 3 and 6 and videos were obtained with a high contrast instrument (Biotek, Cytation 5) for 72 hours.

### Cell viability assay

To measure the cell viability, hMSC spheroids at day 1, 3 and 6 were stained with Calcein-AM (BIOLEGEND, 425201) and PI (SIGMA, P4170-25MG) according to the manufacturer’s protocol. Briefly, Spheroids were incubated in PBS containing Calcein-AM (0.01µM) and PI (3 µM) at 37 °C for 1 h, washed twice with PBS, and then resuspended in MiPS/L-KSR medium.

### Library construction and RNA sequencing in BGISEQ-500 platform

To perform RNA sequencing, Cells were first collected and lysed by Trizol and total RNA was extracted according to the manufacturer’s instructions (Invitrogen, 10296-028). 1 ug of total RNA sample was purified using poly-T oligo-attached magnetic beads, and the purified mRNA was then fragmented using divalent cations under elevated temperature. First strand cDNA was generated using reverse transcriptase and random primers and the second strand cDNA was synthesized using DNA Polymerase I and RNase H. The synthesis product was purified with DNA clean beads, followed by end repair, A-tailing and subsequent ligation to the adapter. After purification and PCR amplication, the products were subjected to single strand circularized DNA molecule (ssDNA circle) preparation for final library construction. DNA nanoballs (DNBs) were generated with the ssDNA circle by rolling circle replication (RCR) to intensify the fluorescent signals during the sequencing process. The DNBs were then loaded into the patterned nanoarrays and sequenced on the BGISEQ-500 platform using pair-end 100 strategy for sequencing and subsequent data analysis [24].

### Initial processing and alignment of RNA-seq data

The FASTQ data of each sample were aligned to the rRNA database (downloaded from NCBI) by SOAPaligner (version 2.21t) to remove rRNAs, and the remaining reads were processed with SOAPnuke (version 1.5.3) [25] to trim adaptors and filter out the low-quality reads. The filtered data were aligned to the hg19 RefSeq transcriptome downloaded from the UCSC Genome Browser database [26] using bowtie2 (version 2.2.5) [27]. Quantification of gene expression levels in raw counts and FPKM for all genes in all samples was performed using RSEM v1.2.4 [28].

### Identification of differentially expressed genes

Differential expression of genes in each group was determined using the R package Deseq2 [29] with default parameters, in which an adjusted p-value less than 0.05 and log2 (fold-change) > 1 was used to identify significantly differentially expressed genes.

### GO term and KEGG enrichment analysis

Gene ontology and KEGG pathway enrichment were analyzed using DAVID [30] and the BH method was used for multiple test correction. GO terms with an FDR less than 0.05 were considered as significantly enriched.

### Flow cytometry

Monolayer MSCs recovered from spheroids at day 6 were harvested and dissociated into single cells by trypsinization and pipetting. To determine cell surface antigen expression, the samples were incubated with the following antibodies: human monoclonal antibodies against CD73 (BIOLEGEND, 344004), CD90 (BIOLEGEND, 328110) and CD105 (BIOLEGEND, 323205). The samples were analyzed using a flow cytometer (BD Biosciences) and gated according by forward scatter and side scatter.

### qPCR

Cells were collected and lysed by Trizol and total RNA was extracted according to the manufacturer’s instructions (Invitrogen, 10296-028). RNA was quantified with a Nanodrop spectrophotometer (Thermo Scientific). 3µg of total RNA was used for reverse transcription with the Prime Script First Strand cDNA Synthesis Kit (Takara, D6110A). Quantitative real time PCR (qPCR) was performed using the TB Green Premix Ex Taq kit (Takara, RR420A). Thermal cycling was performed with a 7500 Real-Time PCR desktop machine (Applied Biosystems) by incubating the reactions at 95°C for 20 s followed by 40 cycles of 95°C for 1 s and 60°C for 20 s. The primers for qPCR analyses are: *STC1* Forward-CACGAGCTGACTTCAACAGGA, Reverse-GGATGTGCGTTTGATGTGGG; *CCL7* Forward-CAGCCAGATGCAATCAATGCC, Reverse-TGGAATCCTGAACCCACTTCT; *HGF* Forward-GCTATCGGGGTAAAGACCTACA, Reverse-CGTAGCGTACCTCTGGATTGC; *IL24* Forward-TTGCCTGGGTTTTACCCTGC, Reverse- AAGGCTTCCCACAGTTTCTGG; *TGFB3* Forward - ACTTGCACCACCTTGGACTTC, Reverse - GGTCATCACCGTTGGCTCA. *GAPDH* Forward - GGAGCGAGATCCCTCCAAAAT, Reverse-GGCTGTTGTCATACTTCTCATGG.

### Data analysis

hMSCs spheroid size was measured with the Image J software. The mean and standard derivation were calculated with excel software.

### Ethical statement

Written informed consent was obtained from donors for all human samples and all experiments were approved by the BGI ethics committee.

## Results

### Human mesenchymal stem cells spontaneously form spheroids in serum-free medium containing KSR

As a substitution of serum, KSR was first used to maintain mouse embryonic stem cells (mESCs). Recently, researchers observed that KSR can promote the proliferation and differentiation of adipose-derived MSCs in monolayer cultures[31, 32], and facilitate the formation of 3D rat testicular culture, indicating that KSR seems to be a suitable substitution of FBS for 2D and 3D cell culture[33]. To test the effect of KSR-containing medium on hMSC culture, we performed a systematical comparison of different hMSC culture medium, including L-FBS, KSR containing MiPS and optimized L-KSR (Figure 1A). Interestingly, when cultured in MiPS medium, a medium originally designed for maintenance and expansion of human embryonic stem cells (hESCs) [34, 35], the dissociated single hMSCs maintained a round cell morphology, attached lightly to the tissue culture dish surface at day 1 and generated spheroids at day 3, while the single hMSCs seeded in L-FBS maintained fibroblast-like morphology (Figure 1B). To further determine the key ingredient(s) in MiPS that promote spheroids formation, we conducted a screening that each time one component was removed from MiPS to establish several incomplete MiPS groups, and test the effect of each group on hMSC spheriods formation. We found that most of these incomplete MiPS groups was sufficient to support spontaneously spheroids formation, except for the group without KSR (Figure 1B), as they attached to the culture dish, maintained fibroblast-like morphology, and mimiced the cells in L-FBS medium (Figure 1B), indicating that KSR is an essential component of MiPS for spheroids formation. To test whether KSR alone was sufficient to generate the hMSC spheroids, we used the basal medium L-DMEM and only added KSR as extra component to culture dissociated hMSCs. Excitingly, we discovered that KSR alone is sufficient for spheroids formation at the concentration as low as 2%, though higher concentration tended to promote a better spheroid formation (Fig S1A-B). Moreover, the addition of KSR alone into other several basal mediums, including RPMI1640, DMEM/F12, H-DMEM, MEM, can also promote the generation of MSC spheroids (Figure S1C), indicating a powerful effect of KSR to substitute serum in these culture medium system for hMSC maintainness. We also tested the effect of initial cell concentration on spheroids formation and found MSCs could generated spheroids at a concentration as low as 1×10^4^/ml in 20% KSR (Fig S2). To simplify, we used 20% KSR in L-DMEM medium in the subsequent experiments and refer to as L-KSR.

**Figure 1.**
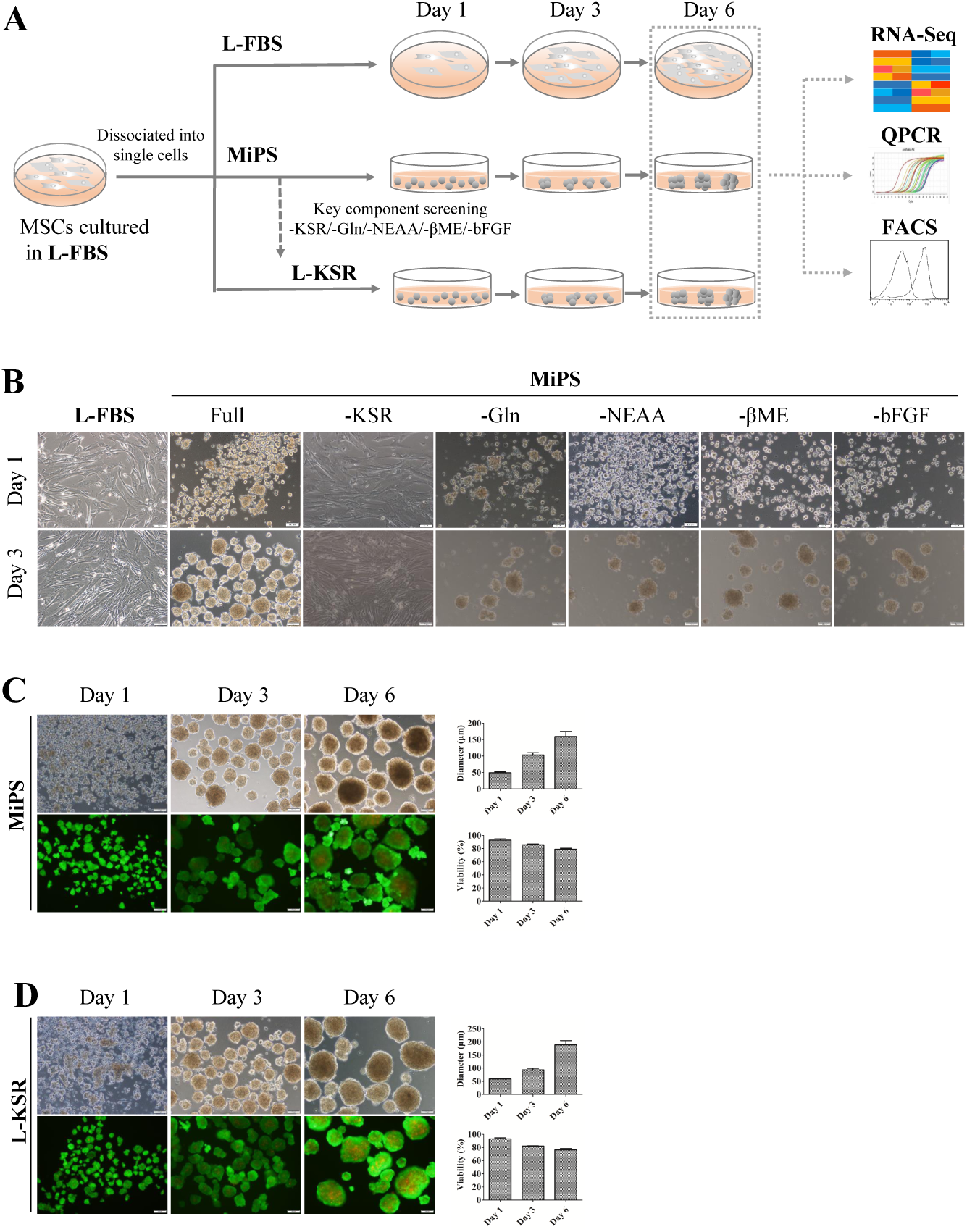
MSCs at P3 can spontaneously form spheroids in medium containing KSR. (**A**) Schematic diagram shows the experimental procedure; **(B**) hMSCs cultured in MiPS and MiPS without KSR/GLn/NEAA/βME/bFGF; **(C-D)** hMSC spheroids was generated and stained with Calcein-AM /PI in MiPS/KSR at day 1, 3 and 6 on tissue culture dishes. Statistical analysis of mean diameter and cell viabilities of hMSC spheroids, sizes were measured from captured images of spheroids (n = 12-20), values are mean ± SD (n = 3), % of live cells (Calcein-AM /PI) was determined using flow cytometry, values are mean ± SD (n = 3). Scale bars:100μm

We then analyzed the hMSC spheroid size and cell viability in L-KSR and MiPS medium system to compare their capability in promoting spheroid formation. In both media, the nearby cells spontaneously migrated and aggregated into small and loose spheroids from day 1 to day 3, and several small spheroids coalesced into large and compact spheroids from day 3 to day 6 (Figure 1C and 1D, supplementary video 1, 2), showing no dramatic diference on the formation speed. In addition, the mean diameter of hMSC spheroids increased with the prolongation of culture time in both media. Cell viability identified by Calcein-AM/PI showed that the percentage of dead cells increased in spheroids in both MiPS and L-KSR with cultured time, but still maintained a high fraction of viable cells at day 6 (>80%), and no significant difference was observed between the two medium groups (Figure 1C and 1D), indicating the effect of L-KSR on hMSC spheroid formation is comparable to MiPS and representing the typical hMSC spheroids phenotype previously reported using a hanging drop protocol [7]. Overall, our data showed that MiPS medium is capable to support MSCs to spontaneously form spheroids, and the medium component KSR is necessary and sufficient to generate spheroids.

### MSCs at high passage retain the ability to form spheroids in KSR medium

Recent studies have shown that MSCs gradually lose their therapeutic potency due to an increasing senescent cell subset during long-term culture *in vitro* [36-39]. To test whether hMSCs at high passage could also form spheroids in KSR medium, hMSCs after 5 or 8 passages in FBS containing medium were cultured in MiPS and L-KSR medium. Our results showed that cells at P5 and P8 could both form spheroids in MiPS/L-KSR medium (Figure 2A and B), although statistical analysis showed that spheroid mean diameter from P8 hMSCs slightly decreased when compared to that of P5 (Figure 2 A and B). Overall, our data demonstrated that MSCs at high passage still can generate spheroids in KSR medium.

**Figure 2.**
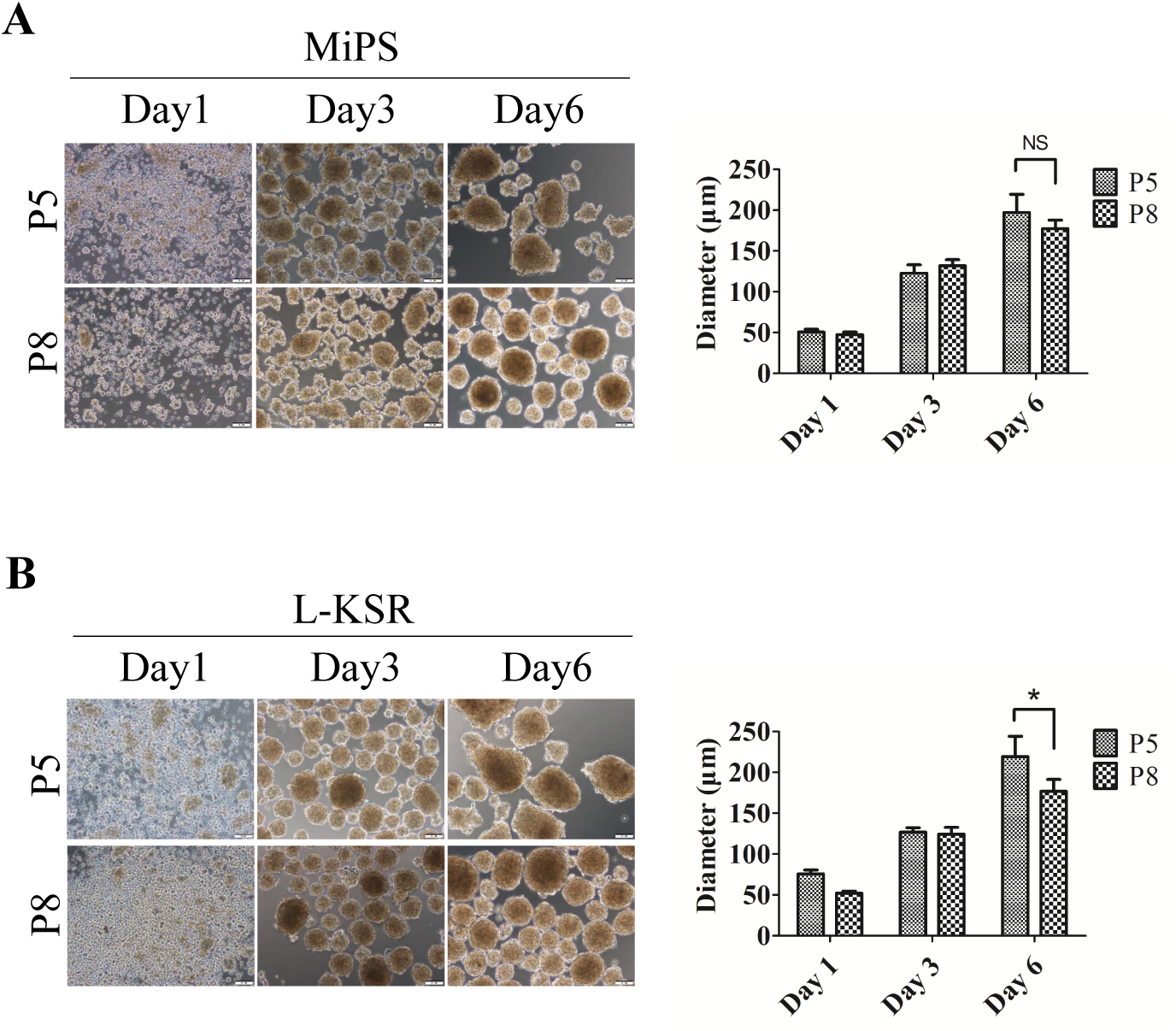
MSCs at higher passage retain the ability to form spheroids in medium containing KSR. **(A)** MSCs at P5 and P8 generated spheroids in MiPS at day 1, 3 and 6 on tissue culture dishes. Statistical analysis of MSC spheroid mean diameter cultured in MiPS; **(B)** MSCs at P5 and P8 generated spheroids in L-KSR at day 1, 3 and 6. Statistical analysis of MSC spheroid mean diameter cultured in KSR, sizes were measured from captured images of spheroids (n = 12-20), values are mean ± SD (n = 3). Not significant (NS) P ≥ 0.05, *P < 0.05. Scale bars:100μm

### Transcriptomics analysis reveals that MSC spheroids generated in KSR medium obtain enhanced expression of therapeutic genes

Previous studies demonstrated the expression of anti-inflammatory factors (*TNFα, TSG-6, STC-1*), and angiogenic growth factors (*ANG, FGF-2, ANGPT-2, HGF*) are significantly increased in MSC spheroid cultures. These results suggested that MSCs cultured in spheroids can enhance cell therapeutic potentials, including anti-inflammation [7] and proangiogenesis [8-10]. To investigate whether the hMSC spheroids generated with our method may acquire similar advantages, we plated P5 and P8 MSCs in L-FBS, MiPS, and L-KSR medium respectively and collected the cells or hMSC spheroids after 6 days in culturing in those different mediums for RNA-seq analysis.

We analyzed the overall transcript expression level in these three groups. Transcriptomic correlation analysis showed that all the samples were highly related (Figure 3A and B), and heatmap results showed that the expression pattern of several important MSC marker genes was similar among the three groups (Figure 3C), demonstrating that these samples maintained MSC features. To validate whether genes of therapeutic potential were altered in 3D spheroids, we compared 3D spheroids in MiPS/L-KSR with 2D adherent monolayer MSCs in L-FBS. The heatmap results showed that potentially therapeutic genes, including *STC1, CCL7, TNFRSF1B, LIF, TGFB, IL1B, IL1A, HGF* were up-regulated in 3D spheroids with MiPS/L-KSR medium, while *DKK1, VIM* genes were down-regulated (Figure 4A). Gene Ontology (GO) enrichment analysis showed that these changed genes are associated with extracellular matrix organization, cell adhesion, wounding healing, angiogenesis, inflammatory response, signal transduction, and immune response (Figure 4B). KEGG analysis showed that these genes are associated with protein digestion and absorption, ECM-cytokine receptor interaction, focal adhesion, and P13K-Akt signaling pathway (Figure 4C). Our qPCR assay confirmed the up-regulated expression of *STC1, CCL7, HGF, IL24*, and *TGFB3* in 3D spheroids with MiPS/L-KSR medium (Figure. 4D), which are considered as critical genes required for MSCs’ function. In summary, the RNA-seq and qPCR results suggested that the expression of potentially therapeutic genes in spheroids can be enhanced in MiPS/L-KSR.

**Figure 3.**
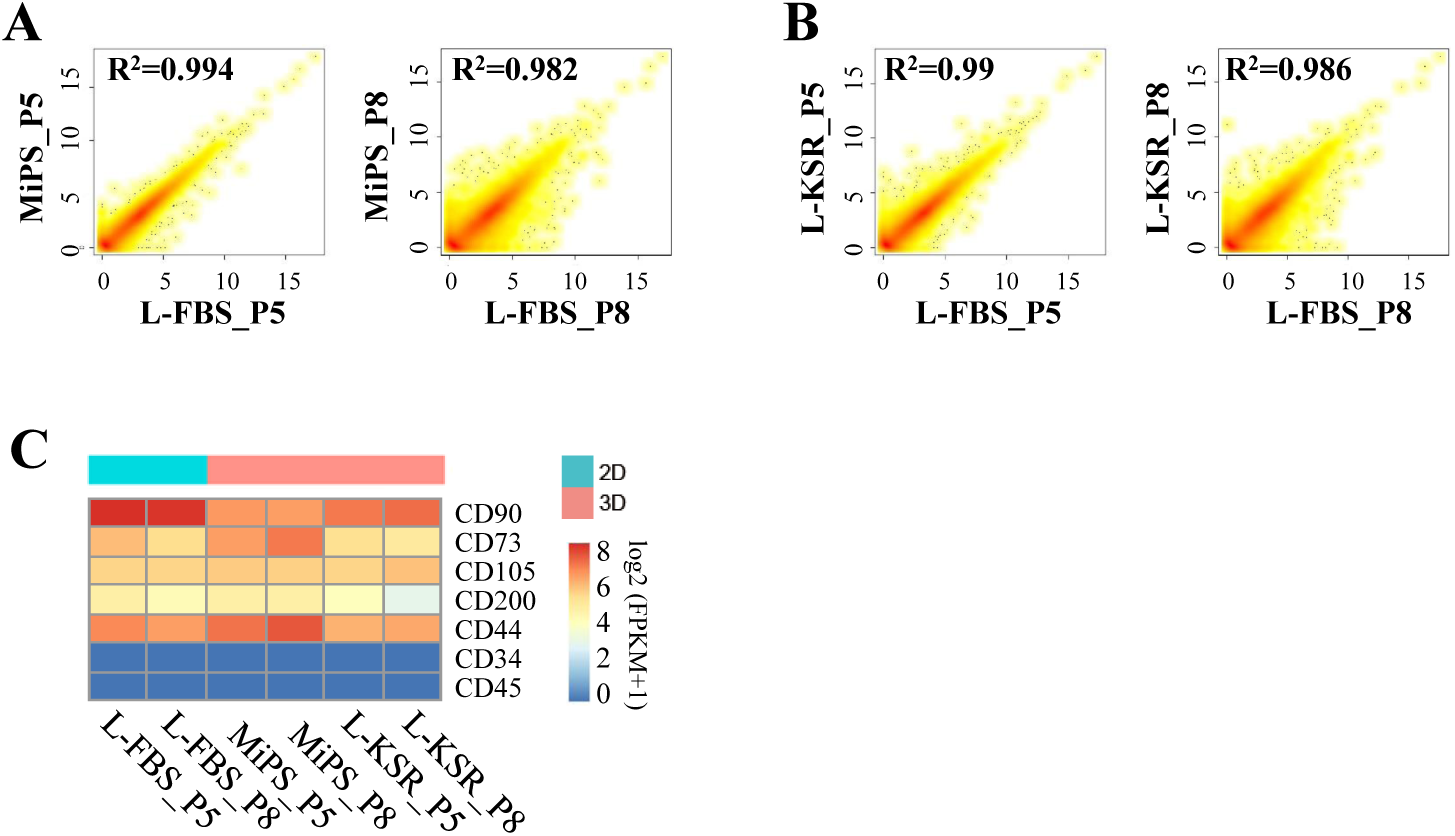
Transcriptomic correlation analysis of spheroids at day 6 at P5 and P8. **(A-B)** Comparison of RNA-Seq gene expression profiles between spheroids in MiPS/L-KSR at P5 and P8 and their corresponding FBS control. R^2^ stands for correlation coefficient; (**C**) Heat map shows the gene expression level of several specific makers of hMSCs.

**Figure 4.**
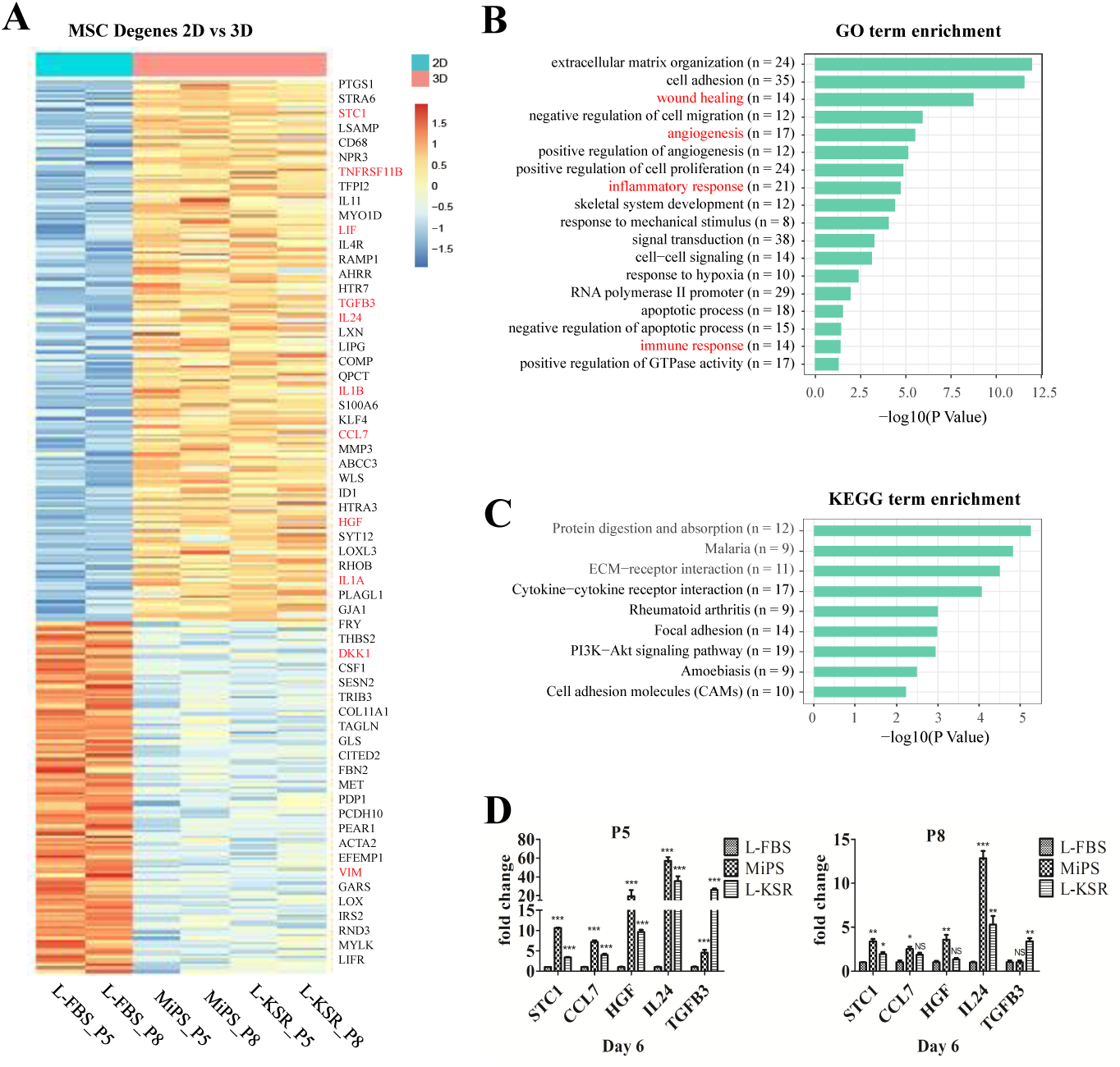
Transcriptomic expression analysis of hMSC spheroids. (**A**) Heat-map shows scaled expression [log2 (FPKM+1)] of discriminative genes between 3D spheroids (both in MiPS and L-KSR) and 2D normal MSCs (in L-FBS medium) at day 6 at P5 and P8. adjust P value < 0.05. Color scheme is based on z-score distribution from – 2 (blue) to 2 (red). **(B)** Gene ontology (GO) analysis between 3D and 2D, “n” indicates gene numbers. **(C)** KEGG analysis between 3D and 2D. **(D)** qPCR results analysis of *STC1, CCL7, HGF, IL24* and *TGFB3* between 3D and 2D cells at P5 and P8 at day 6. The expression levels of mRNA were normalized to *GAPDH*, and expressed as a ratio of mRNA levels of genes of interest to that of *GAPDH*. Not significant (NS) P ≥ 0.05, *P < 0.05, **P < 0.01, and ***P < 0.001.

### Spheroid-recovered MSC cells retain mesenchymal stem cell features

Previous studies showed that the cells in spheroids retained most of the surface epitopes of hMSCs from adherent cultures [4]. To ensure that the MSCs in our KSR culture system retain these MSC properties thus maintaining their value for research and clinical applications, we transferred spheroids derived from P8 MSC at day 6 from MiPS and L-KSR back to standard L-FBS culture medium. Results showed that the MSC spheroids began to attach on to the culture dish surface, and then, from day 1 to day 6, cells in the spheroids migrated out and adhered to culture dish and proliferated (Figure 5A). Spheroid-recovered MSC cells were collected for FACS analysis. FACS results showed that the expression of MSC markers, including CD73, CD90, and CD105 in MiPS/L-KSR samples were still highly expressed, which is similar to that in normal cultured 2D MSCs (Fig. 5B).

**Figure 5.**
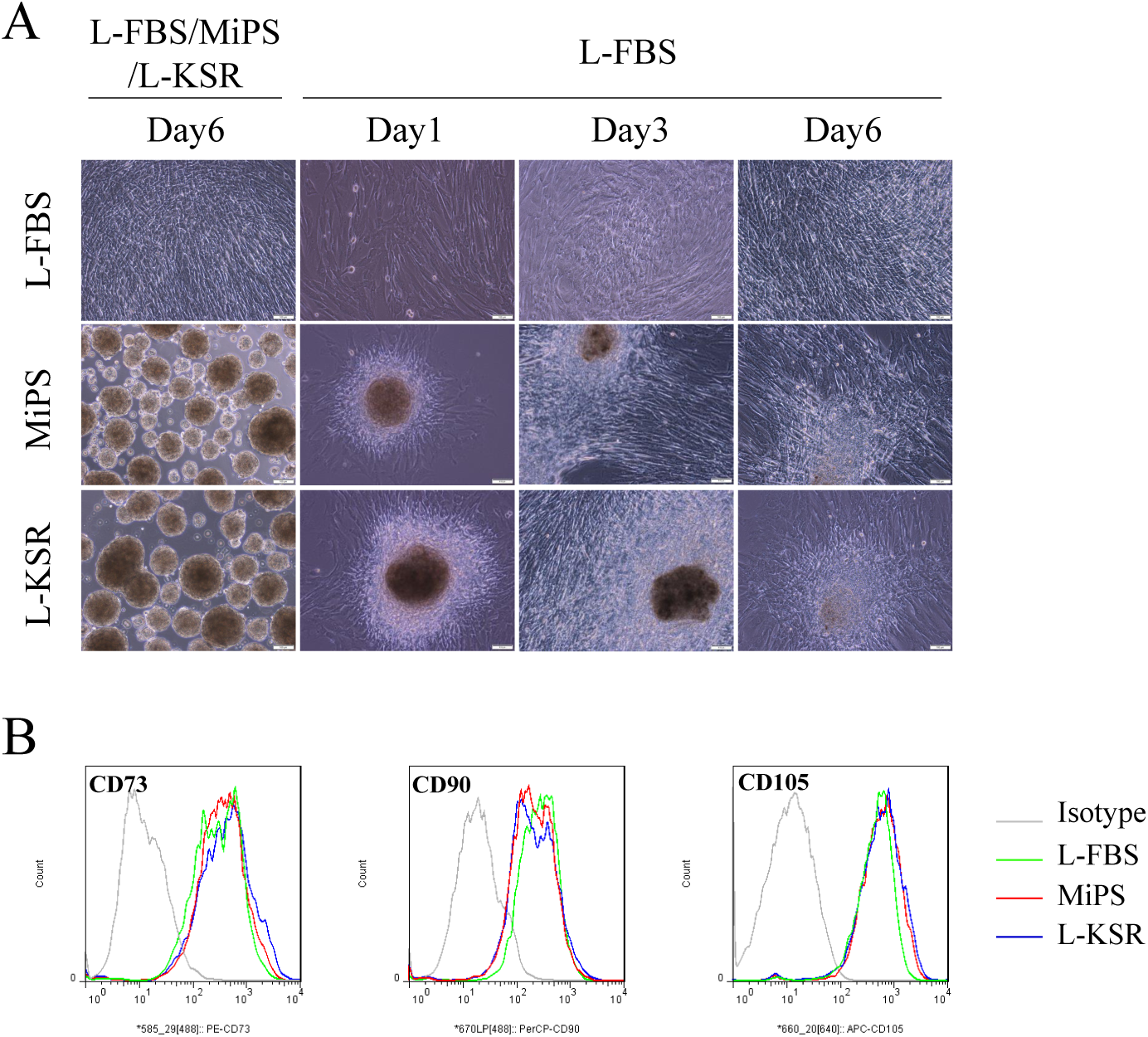
Spheroid recovered hMSCs retains mesenchymal stem cell features. (**A**) Cells in spheroids at day 6 at passage 8 in MiPS/L-KSR migrated out and adhered to tissue culture dishes when reseeded in L-FBS medium. Cells cultured in FBS as a control; **(B)** FACS analysis of MSC positive markers CD73, CD90 and CD105 of spheroid recovered hMSCs in MiPS/L-KSR. Cells cultured in L-FBS as a control. Scale bars:100μm

## Discussion

Cells cultured in 3D as spheroids *in vitro* provide enhanced cell-cell interactions and more closely mimic the natural microenvironment of a tissue. Therefore, it has been widely used in various fields, including tumor biology [40, 41], drug discovery, toxicology screening [42, 43] and organoid research [44]. Previous studies showed that MSCs can form spheroids with a variety of methods. However, most of those methods have to use medium containing FBS or need special instruments (agarose microwells) or complicated procedure (hanging drops or gel coating) [12], hindering their large-scale implication in clinical applications. Here, we developed a convenient method to generate MSC spheroids spontaneously in a novel serum-free formula without any special instrumentation or pre-coated gels.

First of all, we found that MiPS could prevent cells from adhering to the tissue culture dish, and facilitate cells to migrate and aggerate into spheroids (Fig. 1B). Then, we demonstrated that KSR in MiPS was the only critical active ingredients to promote spheroid formation. In fact, KSR is not only necessary, but also sufficient to promote hMSC spheroid formation when added into different basal medium (Fig.1 B-D and Fig. S1). KSR is a substitution of FBS and all the components are well defined, consisting of albumin, transferrin, insulin, collagen precursors, amino acids, vitamins, antioxidants, and trace elements [33]. Therefore, KSR is more suitable for clinical-grade MSC production as it eliminates many of the uncertainties encountered when using poorly defined serum supplements. Our study demonstrated that KSR at a concentration as low as 2% could promote hMSC spheroids formation, though a higher concentration tends to generate larger spheroids in a relatively shorter time (Fig. S1). It would be interesting to further define the key components in KSR that are pivotal to the spheroid’s formation.

Previous studies showed that short-term culture of MSCs in a 3D environment had no significant effect on the level of MSC-specific immunophenotypic marker expression [11]. In this study, we compared the expression pattern of hMSC spheroids derived from MiPS and L-KSR with normal cultured hMSCs through RNA-seq. The overall gene expression pattern, including the MSC marker genes and pluripotency-associated genes, is similar between hMSC spheroids and hMSCs (Fig. 3), suggesting our method didn’t change the basic properties of hMSCs. Various studies have demonstrated MSC cells in spheroid generated in medium containing FBS have a higher expression level of immunomodulatory related factors [7, 45]. However, these mediators were not up-regulated in spheroids cultured in the chemically defined xeno-free medium [20]. Interestingly, we found hMSC spheroids generated in KSR medium up-regulated the expression of potentially therapeutic genes (Fig. 4A and D). Go Term and KEGG analysis of these differentially expressed gene indicated that the signaling pathways enriched in KSR derived hMSC spheroids were associated with extracellular matrix organization, cell adhesion, wounding healing, angiogenesis, inflammatory response, signal transduction, and immune response, which were considered to be related to the therapeutic function of MSCs (Fig. 4B and C). More importantly, when our hMSC spheroids were re-plated to a tissue culture dish in FBS contained medium, spindle shaped cells migrated out and retained MSC properties (Fig. 5). These results suggest that the hMSC spheroids may be implanted directly into the body and served as an MSC reservoir to play a sustained therapeutic role, though the real biological functions of the hMSC spheroids generated with our method need further *in vivo* validation studies.

In conclusion, we developed a practical and convenient method to generate hMSC spheroids in a defined serum-free medium and preliminary studies suggest it enhanced the therapeutic effect of hMSCs. We anticipate the hMSC spheroids generated with our method could be widely used for future clinical research and therapy.

## Supporting information

FiugeS1

## Author contributions

G.D. designed the experiments. G.D., Y.G., Q.C., and Q.D. performed the experiments. S.W. processed the RNA sequencing data. S.W. and G.D.analyzed the data. W.Z and J.T. collected the umbilical cord. Q.W., Z.S., W.O., J.L. and Y.G. jointly the discussions. G.D. and S.W. wrote the manuscript. Y.G. and C.L. revised the manuscript. Y.G supervised the study. All authors reviewed and approved the final manuscript.

## Acknowledgments

We thank Micheal Dean for help editing the language and BGI colleagues who helped to produce the high-quality data.

## Availability of supporting data

The data that support the findings of this study have been deposited in the CNSA (https://db.cngb.org/cnsa/) of CNGBdb with accession code CNP0000456.

