## Supplementary material for "Serum-free culture system for spontaneous human mesenchymal stem cell spheroids formation": FiugeS1

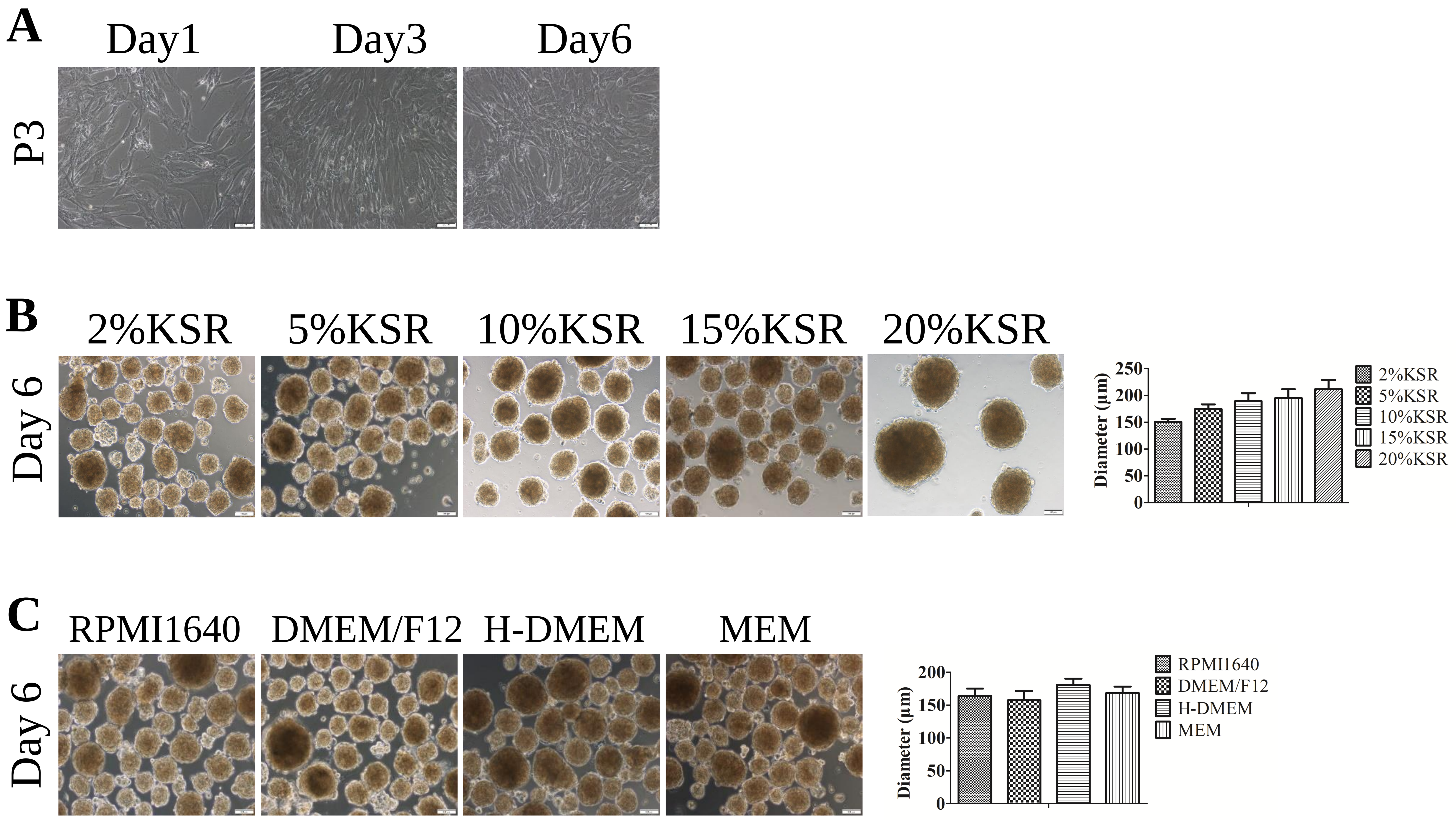


**Figure S1 MSC cells can form spheroids in KSR contained medium.**

**(A)** hMSCs cultured in L-FBS at passage 3; **(B)** hMSC spheroids at day 6 generated from hMSCs at passage 3 in L-DMEM at the indicated concentration KSR. Statistical analysis of hMSC spheroids mean diameter cultured in different concentrations of KSR in L-DMEM medium; **(C)** hMSC spheroids at day 6 generated from hMSCs at passage 3 in various basal medium, including RPMI1640, DMEM/F12, H-DMEM, MEM with 20% KSR. Statistical analysis of hMSC spheroids mean diameter cultured in various basal medium with 20% KSR, sizes were measured from captured images of spheroids (n = 12-20), values are mean ± SD (n = 3). Scale bars:100μm
