## Supplementary material for "Serum-free culture system for spontaneous human mesenchymal stem cell spheroids formation": Figure S2

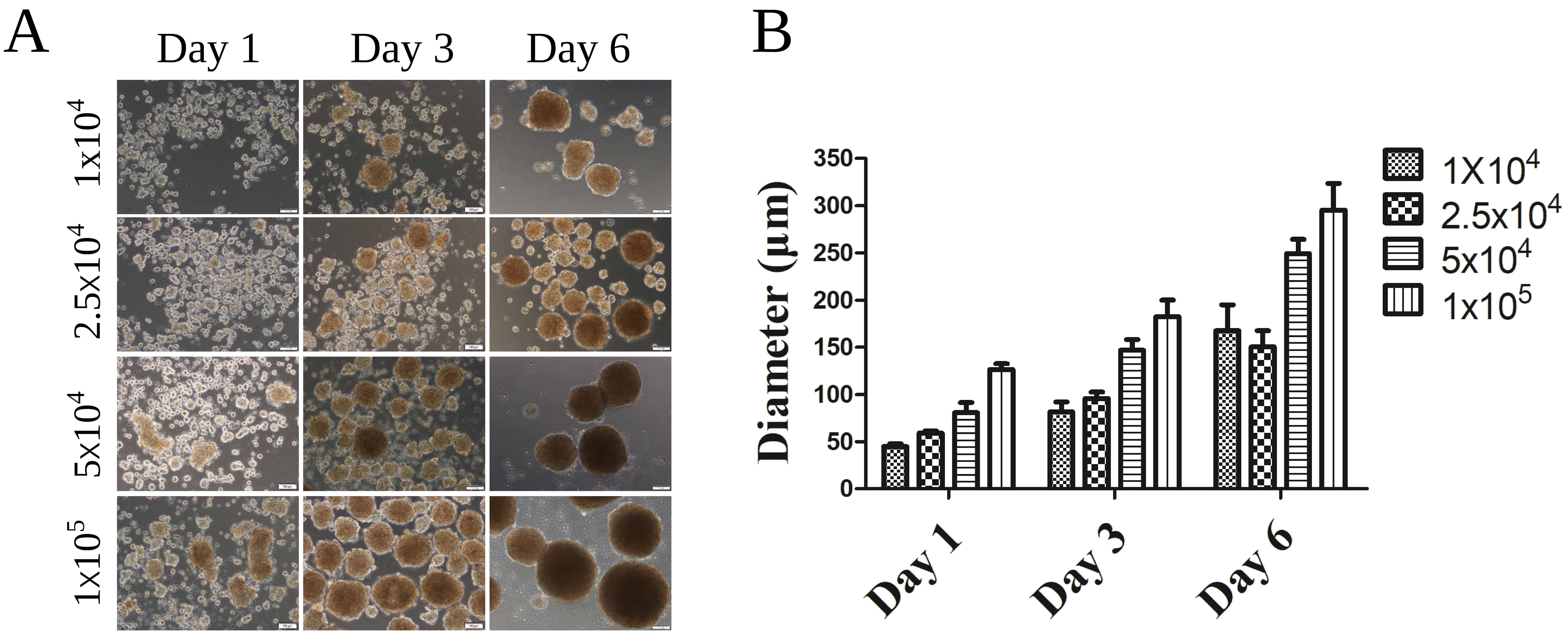


**Figure S2 MSCs can form spheroids at different cell concentrations in 20% KSR containing medium.**

**(A)** hMSC spheroids generated from MSCs at passage 3 in L-DMEM with 20% KSR at different cell concentration; **(B)** Statistic analysis of hMSC spheroids mean diameter cultured in different concentrations of KSR in L-DMEM medium, sizes were measured from captured images of spheroids (n = 12-20)， values are mean ± SD (n = 3). Scale bars: 100μm
